# Differential functional connectivity between hippocampus and prefrontal cortex is associated with heterogeneity in open field exploration

**DOI:** 10.1101/2025.09.22.677917

**Authors:** Robert G.K. Munn, Desiree D. Dickerson, Amy R. Wolff, David K. Bilkey

## Abstract

A common paradigm in behavioral neuroscience involves recording neural activity from freely moving rodents as they forage in an open field. This procedure is often used in studies investigating spatial navigation, where recording is conducted in regions such as the hippocampus and entorhinal cortex. It is usually assumed that there is no systematic variation in behavior in this paradigm, thereby allowing spatial representations to be examined without the confounding effects of behavior change. Here we show that the behavior of rats in this paradigm can be algorithmically divided into at least two distinct modes, and that the transitions between these modes is marked by distinct differences in theta and gamma band power in the hippocampus, as well as transition-associated communication between hippocampus and prefrontal cortex, with information strongly flowing from hippocampus to prefrontal cortex during the transition period. Moreover, we show that following a maternal immune activation intervention, intra-regional changes in power are preserved, but communication between hippocampus and prefrontal cortex is impaired. These findings demonstrate that animals in the open field perform distinct behaviors that are accompanied by marked changes in brain activity and regional communication.

## INTRODUCTION

A paradigm that is commonly used in behavioral neuroscience involves recording neural activity in the brain of freely moving rodents that are foraging for randomly distributed food items in an open field. It is usually assumed that there is no systematic variation in behavior in this paradigm, thereby allowing neuronal representations to be examined without the confounding effects of behavior change. However, organisms that forage for food or other resources have to continuously make decisions about whether it is best to remain in the current region and exploit the resources that are available locally, or leave and explore for new rewards^1^. Whereas exploitation often involves the animal taking tortuous paths in a local-search region, exploration involves more linear and extensive trajectories. These potential switches of behavioral mode, and the brain mechanisms that underlie them, are usually ignored in open field recording sessions.

Previous investigations of the neurobiology of exploit/explore mechanisms have focused on brain regions such as the prefrontal cortex, amygdala and midbrain dopaminergic regions cortex^2–5^. For example, the behavioral flexibility that underlies mode-switching between local search and exploration behavior has been shown to depend on frontal cortical regions^2,6^. Furthermore, these brain areas are also involved in tracking uncertainty^7^ and the decision to forego immediate reward for exploration in search of greater gains^8^. In many cases, however, explore/exploit decisions also require access to the spatial, temporal, and contextual information that allow discrimination between resource-rich areas, and those that have been previously depleted or are dangerous. Since the hippocampus has a critical involvement in encoding, storing, and retrieving the spatial information necessary for navigation and environmental representation^9,10^, it could potentially support the spatial decision-making that would guide explore/exploit decisions. Furthermore, as hippocampal representations are also responsive to reward^11–13^, and time^9,14^ they could also link time-discounted resource availability to spatial position.

There has been relatively little previous focus on the potential role of the hippocampus in explore-exploit foraging decisions, although one recent study has shown that the human posterior hippocampus is involved in the invigoration of exploration behavior, while the anterior hippocampus supports the transition to exploitation^15^. We predict, therefore, that the behavior of animals foraging in the open field can be categorized into separate explore and exploit modes, and that hippocampal activity will be responsive to these mode changes. In particular, we hypothesize that the integrated spatial, temporal, and reward information that is represented in the hippocampus will be available to prefrontal cortex during the explore-exploit mode transitions. We tested this hypothesis in rats foraging for randomly located food items in an open field and observed that the animals regularly changed from the circuitous, area-restricted, local-search paths that are associated with exploitation^16^ to the more linear and extensive trajectories that are associated with exploration^17^. We then examined LFP activity in the hippocampus during these mode switches and described how they relate to simultaneously recorded LFP and single unit neural activity in the prefrontal cortex, with a focus on the changes of synchronization that occurred between the two regions during explore-exploit mode transitions. Finally, we examined these events in animals that had undergone a maternal immune activation (MIA) intervention, a model of a risk factor for schizophrenia and autism^18^ Previous studies have shown that MIA animals have deficits in memory ^19,20^ and changes in hippocampal and prefrontal function^21–25^ that we hypothesized would compromise communication between these regions.

## METHOD

Data obtained during the experiments described in Dickerson et al (2010)^22^ were reanalyzed in the current study.

### Animals

Twelve female Sprague Dawley rats obtained from the University of Otago Animal Breeding Station were mated at 3 months of age. While under halothane anesthesia, six experimental-group dams were injected with a single dose of Poly I:C (4.0 mg/kg, i.v.) dissolved in saline, whereas control group dams received vehicle, on gestational day 15. Pups were culled to give a litter size of six males and weaned on postpartum day 21. The 12 animals used in this experiment (six MIA and six control) were obtained from independent litters and dams. Animals were tested for prepulse inhibition (PPI) from 3 months of age and were at least 5 months of age at the time of electrode implantation for subsequent electrophysiological recordings. No other procedures were conducted during the interim period. Experimental animals were housed three to a cage within their treatment group before surgical procedures and maintained on ad libitum food and water on a 12 h light/dark cycle.

### Surgery and electrode implantation

Animals were subsequently surgically implanted with an adjustable microdrive assembly for neuron recording using techniques we have described previously^26–32^. Briefly, while anesthetized with a ketamine–domitor mixture, a miniature, moveable microdrive containing seven recording electrodes (Formvar-coated, nichrome wires with 25mm diameter; California Fine Wire) was chronically implanted into the mPFC targeting the deep layers of the prelimbic and infralimbic cortices [3.2 mm anteroposterior (AP) and 0.6 mm mediolateral (ML) to bregma; 3.2 mm dorsoventral (DV) from dura] through drilled stereotaxic-guided trephines. Furthermore, one non-moveable 127mm-diameter, nickel-chromium coated wire was implanted in the CA1 region of the dorsal HPC (3.8 mm AP and 2.5 mm ML to bregma; 2.5 mm DV from dura) to record EEG from near the hippocampal fissure. The assembly was fastened to the skull with jeweler’s screws and acrylic dental cement. Animals remained on ad libitum food and water supply until 2 weeks after surgery, at which time their food was reduced to maintain the animal at 85% of their free-feeding weight to optimize behavior during the recording procedure.

### Electrophysiological recordings

Extracellular spikes were recorded using the dacqUSB multichannel recording system (Axona Ltd.). Signals were buffered via a field effect transistor source-follower mounted to the head stage with a “quiet” electrode as an indifferent. Neuron activity was bandpass filtered (between 360 Hz and 7 kHz), amplified 100 times, and digitized at a 48 kHz sampling rate by the dacqUSB system. Neuronal signals were digitized when the spike on any channel exceeded a threshold set above the background noise levels and were then stored for offline analysis. EEG signals were referenced to a skull screw located anterior to bregma and contralateral to the mPFC electrode trephine, low-pass filtered at 500 Hz, and sampled at 4800 Hz.

### Recording apparatus and procedure

The experimental chamber was a black, plastic, circular tub with a floor diameter of 74 cm and a wall height of 56 cm. The room was lit by a low-level light source located at one side of the room. This was not directly visible to the animal while in the apparatus. Animals were handled for 3 d before experimentation and habituated for two 20-min sessions within the environment for 2 d before their first recording. Two 10-min recordings were then obtained each day (morning and evening) while animals foraged for chocolate hail in the familiar open-field environment. Small quantities of chocolate hail were scattered throughout the chamber at varying intervals to ensure there were always some quantity of food available. The electrode was advanced through the mPFC cell layer by ∼40microns after each recording session.

Single-neuron activity was manually identified on the basis of spike waveform characteristics using an offline cluster cutting program (TINT; Axona Ltd.). Raw EEG, spike times, and positional and behavioral data were read into MATLAB (v2025a, MathWorks) for additional analysis.

The phase of hippocampal theta activity was determined by bandpass filtering the data between 6 and 10Hz using a two-pole Butterworth filter with the `filtfilt` function (MATLAB 2025a; MathWorks) to avoid phase shifts caused by the filtering process and then applying the standard Hilbert transform. Phase zero was the positive peak of the waveform recorded from around the CA1 fissure. To determine the strength of phase-locking and preferred phase of firing of mPFC neurons to underlying LFP activity, each spike was given a phase of firing value with reference to the simultaneously recorded LFP and a measure of angular dispersion (mean vector length - r) and mean phase angle were calculated across all spikes. The value of r provides a measure of the concentration of firing about the unit circle and varies inversely with the degree of dispersion within the data such that 0 indicates that firing is so widely distributed that a mean angle cannot be described) to 1, when all firing is concentrated at the same phase angle.

The distribution of mPFC neuronal firing to hippocampal and LFP was assessed using the Rayleigh’s test for circular uniformity^33^ to determine whether the phase of firing distribution was evenly (and therefore without significant mean direction) or unevenly distributed (and therefore possessing a significant mean direction). To compare phase of firing before and after a behaviour mode transition, firing phase was determined for a 500 ms block of data immediately before and after the mode change. Watson’s U^2^ test for two samples of circular data^34^ was used to determine whether there were significant changes in phase angle across the transition point.

### Behavior

The position of the animals’ head was monitored by a ceiling-mounted video camera connected to a tracking system that monitored infrared light-emitting diodes mounted on the head stage. Head position was sampled at 50 Hz, and positional information was made available to the dacqUSB system. Subsequent offline analysis separated the animal’s behavior into period of local search and exploration. Local search was defined as periods where the animal’s trajectory became tortuous. This was detected automatically by a bespoke MATLAB routine. This routine down-sampled position data to 10Hz and then systematically scanned through that path data, looking two seconds ahead of every sample. For each block of data, the speed of the animals and the eccentricity of the path was calculated. The latter was determined by fitting an ellipse using MATLAB’s singular value decomposition (`svd`) function which returned the length of the major (a) and minor (b) axes. The eccentricity was calculated as 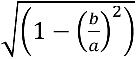 and the range as (*a* + *b*). Two second long path segments where the speed was less than 20cm/sec and range was less than 25cm were labelled as being tortuous. Speed threshold was deliberately set quite high as trial and error revealed that range values were the best detectors of path tortuosity. Adjacent regions of tortuous path that were less than one second apart were joined into a single event. These were regarded as areas of local search, while path trajectories outside these regions were regarded as exploration.

We calculated descriptive statistics for time spent in, and between, local search and also examined the tendency for animals to return to near a region that had been searched already. This involved setting a 20cm border around the center of every local search region and then measuring the time, starting one second after the termination of a local search event, until the animal reentered this border region. This was determined for every local search event.

To compare the pattern of local search events within groups we used a method Scanmatch^35^ which is based on the Needleman-Wunsch algorithm used in bioinformatics to compare DNA sequences. Scanmatch itself was developed to compare a series of eye movement fixations and captures similarities between two 2D patterns in both the spatial and temporal domain^36^. Previous studies have shown that visual search shares several of the properties of foraging behaviour^37^. Here we effectively compressed animal paths into a series of local searches, treated as if they were fixations at a mean location, while explorations were treated as the saccades that link them. A Scanmatch score of 1 would indicate a perfect match between two search patterns while zero would indicate no similarities. To analyze the self-similarity of paths within each group (MIA/control), each local-search path conducted in one recording was compared to a randomly selected path from a recording made from another animal in the same group using Scanmatch. This procedure was repeated 1000 times to generate a distribution of path similarities.

### Histological procedures

After study completion, animals were deeply anesthetized with halothane and underwent transcardial perfusion. After fixation, 60micron coronal sections were mounted and stained with thionin, and subsequent examination of these sections was used to verify the placement of the electrodes.

All procedures and experiments were performed in accordance with ethical guidelines of the University of Otago Ethics Committee.

### Data Analysis

#### Spectral Power

Eight-second-long windows of LFP sampled at 4.8kHz were captured centered on each behavioral transition point. For each transition, comparison eight-second-long segments of LFP were captured in the eight seconds immediately prior to the transition windows. Power spectra were background-subtracted by dividing the power spectrum matrix by the mean power in a randomly selected 8-second window centered between 8 and 10 seconds prior to the transition. For non-transition windows, this meant they were background subtracted against the mean power in another non-transition window 8 to 10 seconds prior. Data was processed using MATLAB (v. 2025a, Mathworks, MA) using a combination of custom-written scripts and functions built in to base MATLAB and the Signal Processing and Circular Statistics toolboxes. Power was computed using a continuous wavelet transform (MATLAB function `cwt`) using a morse wavelet with symmetry of 3 and a time-bandwidth parameter of 60.

#### Coherence

Imaginary coherence was calculated as the magnitude-squared wavelet coherence using the function `wcohere` with 12 voices per octave from the Signal Processing toolbox. Coherence was baseline-subtracted from a control non-transition segment of matching two-second length between six and eight seconds prior to behavioral transition

#### Partial Directed Coherence

We calculated short-time partial directed coherence and directed transfer functions using the WOSSPA toolbox^38^.

#### Statistical Analysis

Data were analyzed using a combination of MATLAB (2025a) and Prism (v 10.1.1). Power data was considered on a per-second basis rather than a per-sample basis. To achieve this, data were averaged in second-long windows, giving eight segments. For coherence data, this averaging was done on a 50ms window. Repeated-measures 2-way Analysis of variance was conducted to determine interactions between transition type and time, and then Bonferroni-corrected post-hoc t-tests were used to determine where in the time series differences in power between transition types occurred. Circular statistics (Rayleigh’s z, Watson’s U) were calculated using functions from the `circstat` toolbox^39^

## RESULTS

### Animal Behavior

As described previously^22^, analysis of pre-pulse inhibition (PPI) data confirmed a significant reduction in percentage PPI in the MIA animals (*t*(9) = −3.241, *p* = 0.01) compared to control animals. All animals made many transitions between high and low tortuosity modes throughout each open field recording session. We refer hereafter to epochs of detected low tortuosity as “explore” epochs, and high tortuosity segments as “local search” epochs, in which animals may exploit local resources. Example 8-second-long transition segments time-centered on the transition point are illustrated in Figures 1A and B. The overall pattern of behavior was similar in both MIA and control animals with control animals producing 39 ± 1.4 (SEM) local search events per ten-minute recording while MIA animals generated 38 + 0.9. The duration of these events was also similar (control 11.7 ± 1.1 seconds; MIA 11.0 ± 0.5 seconds) as was the time between events (control 5.0 ± 0.3 seconds; MIA 5.3 ± 0.2 seconds). The position of the local search events, relative to the center of the arena, was also similar across groups (control 30.3 ± 0.4cm, MIA 29.6 ± 0.4 cm).

**Figure 1.**
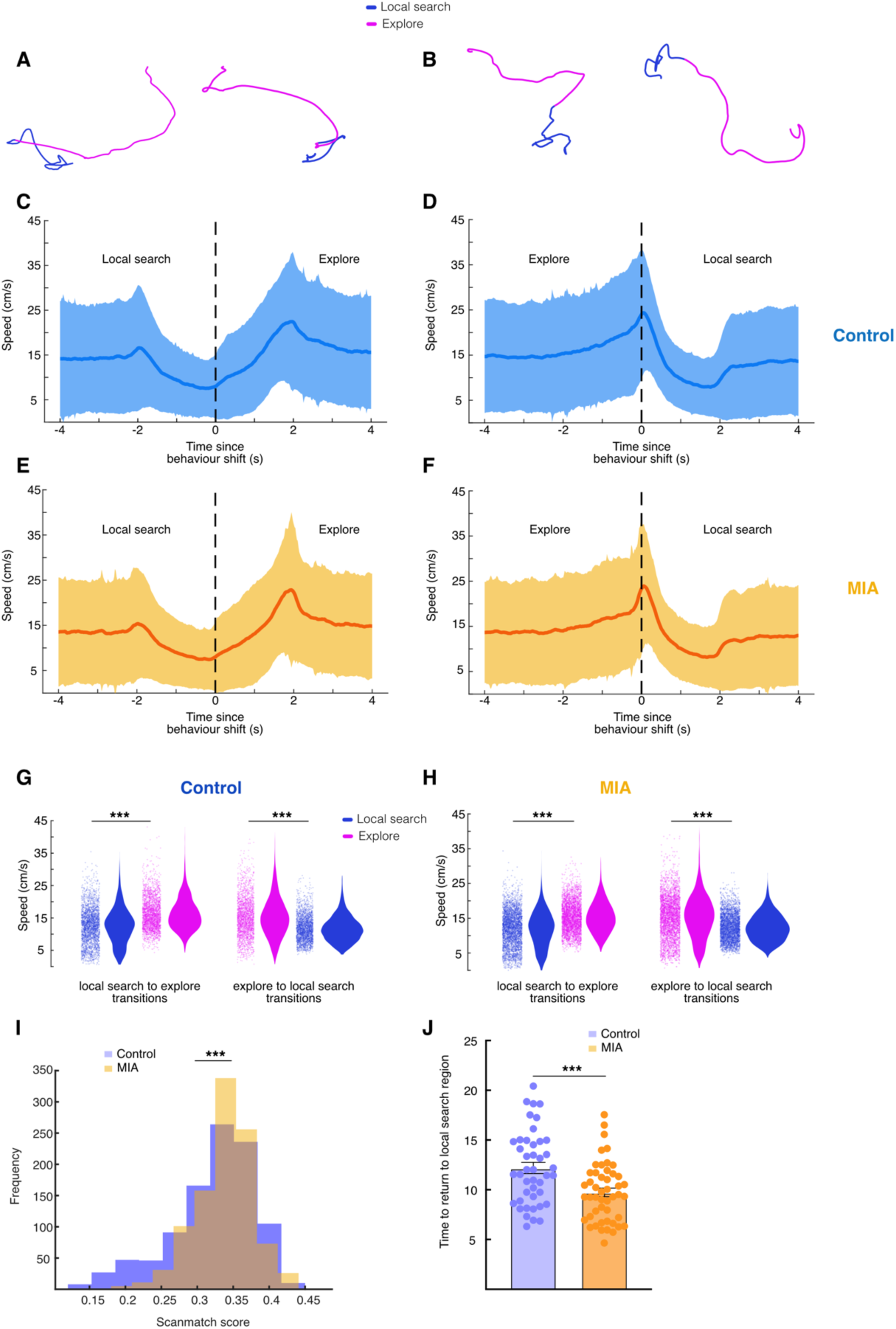
Behavioral changes. **A)** and **B)** each show two eight-second examples of behavioral trajectories when state changes were detected. Four seconds either side of the behavioral change is shown. (A) illustrates high to low transitions, while (B) illustrates low to high transitions. **C)** and **D)** Illustrate the mean speed (solid lines) ± SD (shaded area) from four seconds prior to behavioral state change to four seconds after for captured epochs from control animals. Vertical black dashed lines illustrate the detected behavioral change point. C) Shows speed during high to low transitions, while D) shows low to high transitions. **E)** and **F)** As for (C) and (D), but for epochs captured from MIA animals. **G)** and **H)** show mean speed pre and post behavior shift for both high-to-low (left pair) and low-to-high (right pair) transitions. (G) shows this data for control animals, while (H) shows this data for MIA animals. **(I)** Shows the frequency distribution of scanmatch scores in control (blue) and MIA (orange) animals for 1000 shuffled comparisons across paths. Control animals had a broader, left-skewed distribution of scores compared to the MIA group. **(J)** Illustrates the average time taken for animals to return to a region they had previously searched in each session. Control (blue) animals took longer on average to return to a previously explored local-search area than MIA animals (orange). (***p < 0.001).

As expected, control animals (n = 5) ran faster over all recording sessions (n = 42) during explore behavioral epochs than during local search epochs. During explore-to-local search transitions (n = 2498 transitions), running speed was higher during explore than local search (n = 2497 transitions), (Figure 1D,G explore speed (cm/s) ± SEM = 16.4 ± 3.3 cm/s, local search speed ± SEM = 12.8 ± 1.3 cm/s. t(41) = 10.05, p = 1.27^-12^). The same pattern was true during local search-to-explore transitions (Figure 1C,G local search speed (cm/s) ± SEM = 12.5 ± 2.0, explore speed (cm/s) ± SEM = 16.1 ± 2.4, t(42) = 27.05, p = 9.21^-28^).

MIA animals (n = 6 animals, n = 54 sessions) exhibited the same pattern of behavior, running faster during explore epochs than local search epochs during both explore-to-local search transitions (n = 3228 transitions, Figure 1,E,H; local search speed ± SEM = 11.79 ± 1.25, “low” tortuosity speed = 15.31 ± 1.38, t(53) = 31.7, p = 3.83^-36^) and local search-to-explore transitions (n = 3229 transitions, Figure 1F,H; explore speed (cm/s) ± SEM = 15.19 ± 1.81, local search speed ± SEM = 12.37 ± 0.96, t(53) = 18.17, p = 2.00^-24^).

While overall foraging strategies were similar in both MIA and control animals, an analysis of the self-similarity of paths using the Scanmatch procedure revealed that the distribution of path similarity scores was skewed to the left in control animals (figure 1I). This resulted in significant differences between the control and MIA distribution in terms of both the mean (control 0.322 <u>±</u> 0.002, MIA 0.338 <u>±</u> 0.001; t(1998)= 7.08, p < 0.0001) the shape of the distribution of all comparisons (binned into ten bins; chi^2^ = 94.6, p<0.0001), but also the skew (control -0.88 <u>±</u> 0.07; MIA -0.42 <u>±</u> 0.09, t(43)=4.15, p < 0.0001) of the individual distributions generated by comparing all paths with one other randomly chosen path from the same group. The skew to the left in the control animals’ distribution indicates that there was a greater diversity of search patterns (more lower similarity scores) in control animals, compared to their MIA counterparts. An analysis of the time to return into a region of local search also revealed differences (Figure 1J). MIA animals returned to a region that had been previously subjected to a local search significantly sooner (9.8 ± 0.4 seconds) than control animals (12.2 <u>±</u> 0.6 seconds t(89)=3.515, p =0.0004).

### Spectral power in hippocampus and prefrontal cortex changes with behavior and transition between behavioral states

#### Control animals

##### Hippocampal LFP

Power time series data was analyzed from n = 42 sessions. In the theta (5-12Hz) band, there was a time x transition interaction between explore-to-search and search-to-explore behavioral transitions (F(7,574) = 35.98, p < 0.0001). Bonferroni-corrected post-hoc student’s t-tests revealed that during the second just prior to and ending at the transition point there was more theta power in explore-to-search transitions than search-to-explore transitions (t(82) = 5.30, t < 0.001). One second after to two seconds after the transition there was instead more theta power in the search-to-explore transition segments (t(82) = 7.72, p < 0.001), continuing in the next second (two to three seconds after transition, t(82) = 4.14, p = 0.007). In the beta (13-30Hz) band, there was again an overall time x transition interaction (F(7,574) = 22.5, p < 0.001). In comparison to theta, beta power exhibits the opposite pattern. In the second leading up to the transition point, and for the following second there was more power in the search-to-explore transition segments than explore-to-search segments (-1 to 0 seconds, t(82) = 2.82, p = 0.049, 0 to 1 seconds t(82) = 5.15, p < 0.001). From two to three seconds post transition this pattern reversed, and there was more power in the explore-to-search transitions than search-to-explore (t(82) = 5.81, p < 0.001).

##### Prefrontal LFP

There was no significant interaction or main effect of transition type when comparing between search-to-explore and explore-to-search transitions (Figure 2I, time x transition, F(7,574) = 0.37, p = 0.92, n.s.). By contrast to theta, there was a strong time x transition interaction in gamma power (Figure 2J, F(7, 574) = 6.64, p < 0.0001). Second-by-second post-hoc analysis demonstrated significantly greater gamma power in the second immediately after the transition point in search-to-explore transitions compared to explore-to-search transitions.

**Figure 2.**
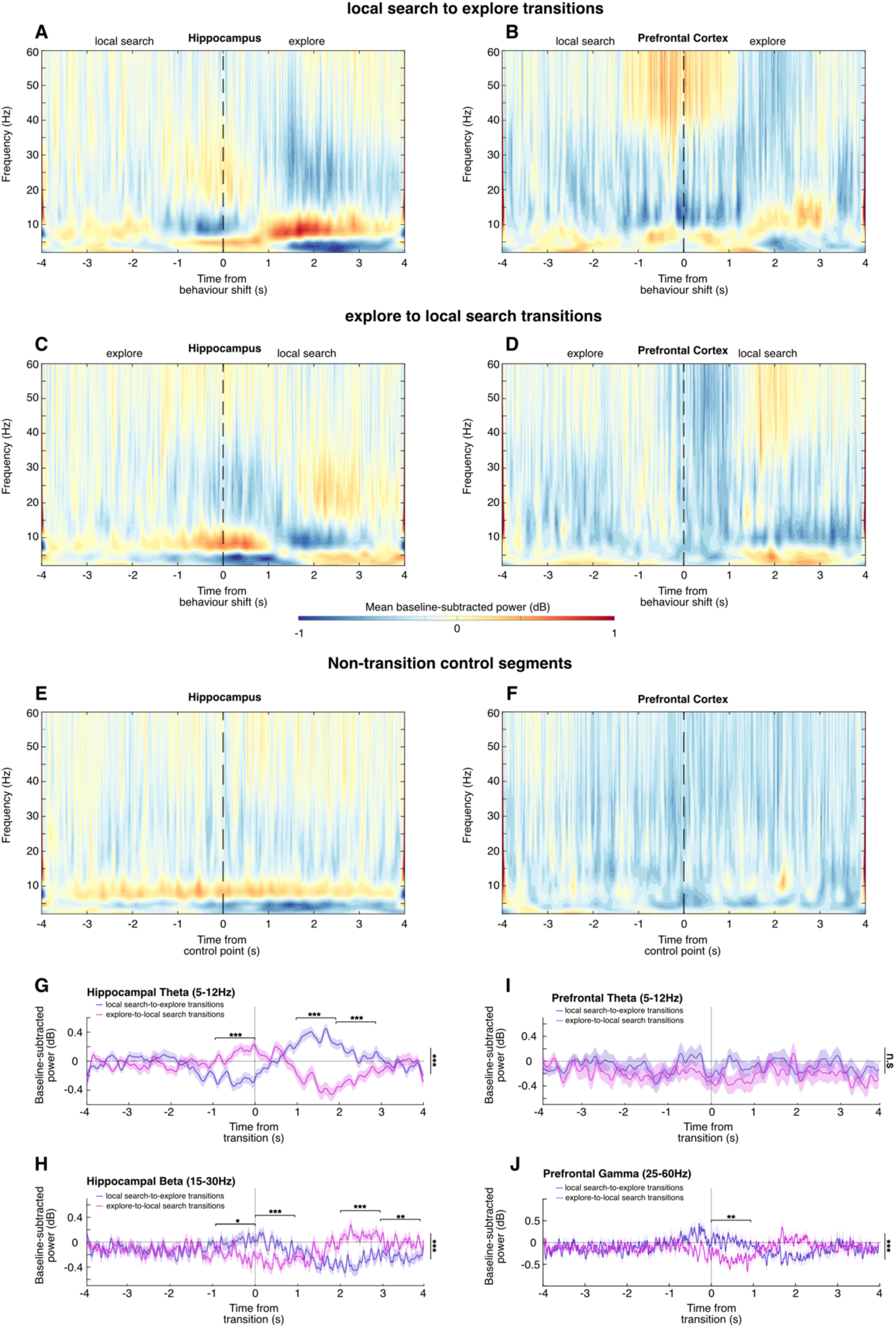
LFP Spectral power in control animals is associated with transitions in behavior. **A)** and **B)** Mean Spectral power from 0-60Hz during the 8 seconds surrounding each exploit-to-search tortuosity transition point. (A) Shows hippocampal LFP and (B) shows prefrontal LFP. **C)** and **D)** as in (A) and (B), but during search-to-exploit transitions. **E)** and **F)** as in (A) and (B), but for non-transition control segments of LFP. **(G)** mean power (solid line) ± SEM (shaded area) in the theta (5-12Hz) band in hippocampus during exploit-to-search (purple) and search-to-exploit (pink) transitions. Horizontal dotted line shows 0 dB, vertical dotted line shows the transition point. **H)** as in (G), but for gamma (25-60Hz) power in prefrontal cortex. **(I,J)** as in (G,H), but for theta and gamma power in prefrontal cortex, respectively. (***p < 0.001, **p < 0.01,*p < 0.05).

#### MIA animals

##### Hippocampal theta

LFP power in the hippocampus of MIA animals followed the same general trend as Control animals. Considering theta-band power, there was an overall time x transition interaction (Figure 3G, F(7,742) = 24.9, p < 0.001). Post-hoc comparisons showed that in contrast to control animals, there was no difference in power between the two types of transition in the second prior to the behavioral transition (t(106) = 2.33, p = 0.175, n.s.). A difference between transition types was only observed between one and two seconds, with significantly more power during the search-to-explore transitions than explore-to-search transitions (t(106) = 8.876, p < 0.001). This pattern was also observed between two- and three-seconds post-transition (t(106) = 4.035, p = 0.0009).

**Figure 3.**
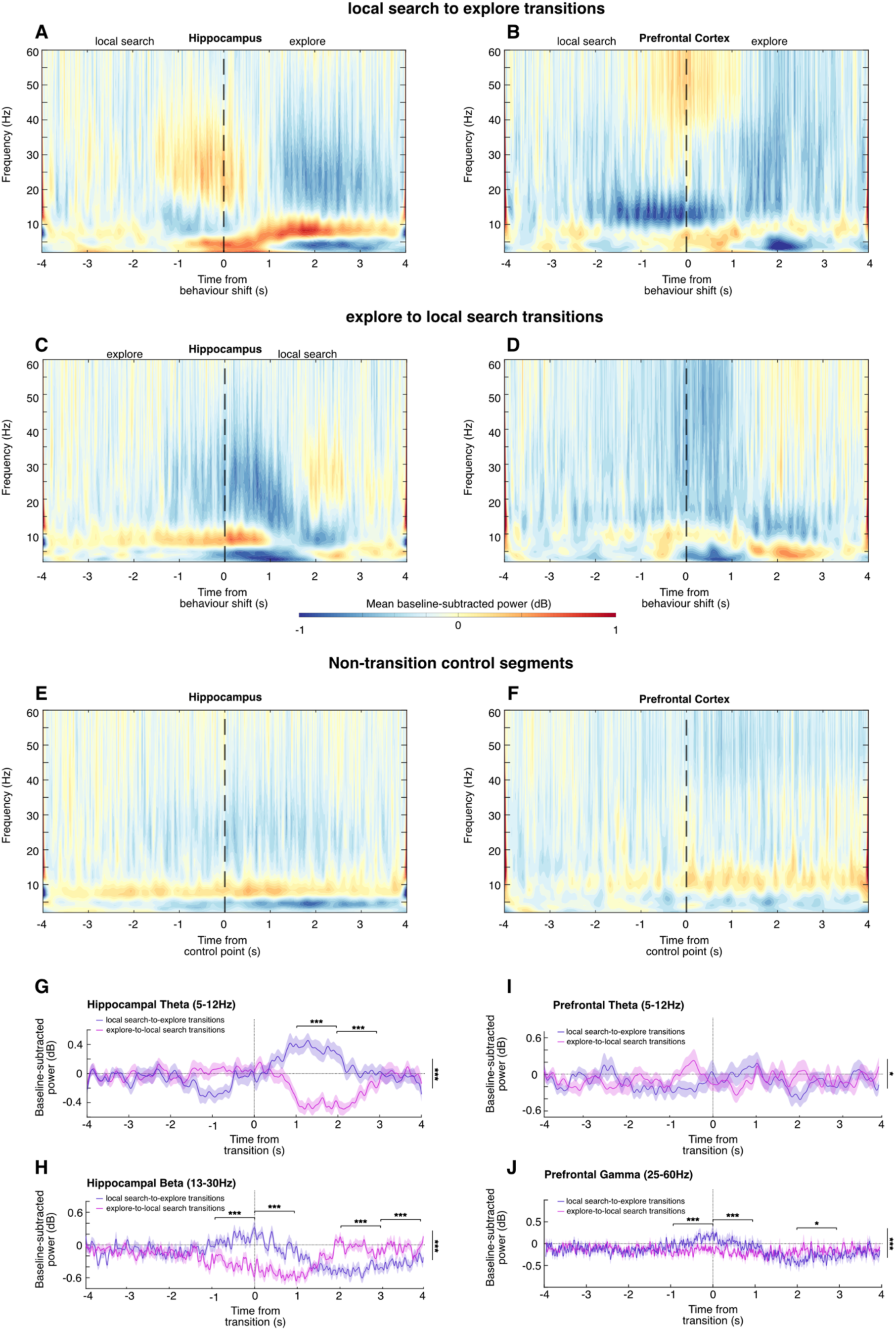
LFP Spectral power in MIA animals is associated with transitions in behavior. **A)** and **B)** Mean Spectral power from 0-60Hz during the 8 seconds surrounding each exploit-to-search tortuosity transition point. (A) Shows hippocampal LFP and (B) shows prefrontal LFP. **C)** and **D)** as in (A) and (B), but during search-to-exploit transitions. **E)** and **F)** as in (A) and (B), but for non-transition control segments of LFP. **(G)** mean power (solid line) ± SEM (shaded area) in the theta (5-12Hz) band in hippocampus during exploit-to-search (purple) and search-to-exploit (pink) transitions. Horizontal dotted line shows 0 dB, vertical dotted line shows the transition point. **H)** as in (G), but for gamma (25-60Hz) power in prefrontal cortex. **(I,J)** as in (G,H), but for theta and gamma power in prefrontal cortex, respectively. (***p < 0.001,*p < 0.05).

##### Hippocampal beta

As with control animals, there was a time x transition interaction in beta power in MIA animals. (Figure 3H, F(7,742) = 41.04, p < 0.001). Post-hoc comparisons showed significantly greater power in search-to-explore transitions than explore-to-search transitions in the second prior (t(106) = 5.71, p < 0.001) and immediately following (t(106) = 6.62, p < 0.001) the transition point. Starting two seconds after the transition, this pattern was reversed with greater power in explore-to-search transitions (2-3 seconds, t(106) = 6.412, p < 0.001; 3-4 seconds, t(106) = 4.07, p = 0.0007).

##### Prefrontal theta

Unlike control animals, MIA animals showed a time x transition interaction in prefrontal theta power (Figure 3I, F(7,742) = 2.32, p < 0.03), although post-hoc analyses revealed no individual differences on a second-by-second basis.

##### Prefrontal gamma

As observed in control animals, MIA animals showed a significant time x transition type interaction in prefrontal gamma power (Figure 3J, F(7,742) = 24.66, p < 0.001). There was significantly more gamma power during search-to-explore transitions than explore-to-search transitions in the second prior to (t(106) = 4.33, p = 0.003) and immediately after (t(106) = 4.11, p = 0.0007) the transition point. This pattern then inverted, with more power in explore-to-search transitions two to three seconds after the transition point (t(106) = 3.05, p = 0.024).

### LFP coherence between hippocampus and prefrontal cortex is associated with behavioral transitions in control but not MIA animals

#### Control animals

Theta band coherence in control animals appeared to be strongly modulated by type of behavioral transition and temporally organized to peak just prior to, and just after, behavioral transition. Overall, there seemed to be greater theta coherence in explore to local search transitions than local search to explore transitions. Analysis revealed that there was a time x transition type interaction (Figure 4E, F(40,160) = 1.58, p = 0.013), as well as a main effect of transition type (F(1,41) = 12.53, p = 0.001). Post-hoc analysis showed that starting 50ms after the behavioral transition point there was significantly greater coherence between hippocampus and prefrontal cortex in the explore-to-search segments (t(83) = 3.25, p = 0.047). There was significantly more coherence in these segments until 350ms after the transition.

**Figure 4.**
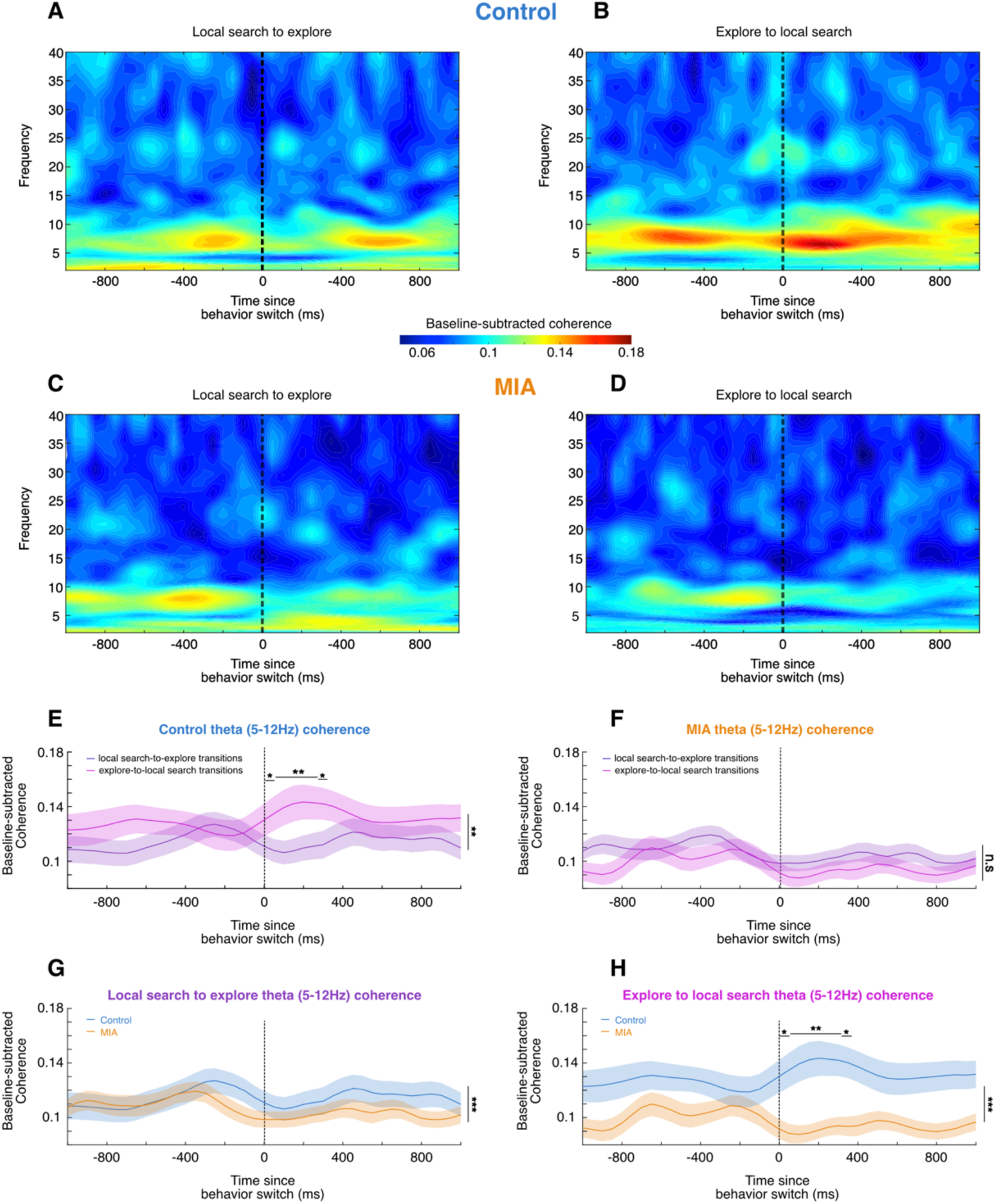
Coherence between prefrontal cortex and hippocampus is associated with behavioral transition type in control but not MIA animals. **(A,B)** Spectral coherence between hippocampus and prefrontal cortex in control animals between 0-40Hz during a two-second peri-transition window. The vertical dotted line indicates the time that the behavioral transition was detected. Local search to explore transitions are shown in (A), and explore to local search transitions are shown in (B). **(C,D)** As in (A,B), but for MIA animals. **(E,F)** Mean (solid lines) ± SEM (shaded area) coherence in the theta (5-12Hz) band during local search to explore transitions (purple) and explore to local search transitions (pink). The vertical dotted line illustrates the time the transition was detected. Control data is shown in (E), while MIA data is shown in (F). **(G,H)** Comparison between mean (solid lines) ± SEM (shaded area) spectral coherence in the theta band in control (blue) and MIA (orange) animals. (G) shows local search to explore transitions, (H) shows explore to local search transitions. (***p < 0.001, **p < 0.01, *p < 0.05).

#### MIA animals

In contrast to the strongly differentiated coherence between transition types, there appeared to be little, if any, modulation of coherence based on types of transition in MIA animals. There was no difference in theta coherence in MIA animals throughout the transition window between transition types (Figure 4C,D,F time x transition F(40,2120) = 0.57, p = 0.986, n.s.). There also appeared to be lower coherence in MIA animals overall. During search-to-explore transitions there was a main effect of animal group, demonstrating overall greater coherence in control recordings than MIA recordings (Figure 4G, F(1,3854) = 21.16, p < 0.001). Comparing MIA and control during explore to local search transitions, there was a main effect of animal group (Figure 4H, F(1,3854) = 260.7, p < 0.001) again revealing significantly higher theta band coherence in control animals compared to MIA. Post-hoc analysis revealed that this effect was significant from 50ms post transition to 400ms post-transition.

### Prefrontal single units phase cluster during transitions

#### Firing rate

A comparison of the mean firing rate of mPFC neurons between the one second pre and one second post the behavioral transition from exploration to local search revealed no change in either the control or MIA group (paired t-tests; control t(104) = 1; p = 0.314; MIA t(106) = 1.5; p = 0.145).

#### Firing phase

An analysis of the phase of firing of mPFC neurons revealed that in both MIA and control animals firing tended to cluster to a particular hippocampal theta phase during the period immediately following the behavioral transition from exploration to local search. The largest vector length was measured near this time point (figure 5A and F) and Rayleigh z tests indicated significant clustering during this transition period (figure 5B and G). A comparison of the phase of firing during the 500ms prior to, and 500ms after, the behavior transition revealed a phase shift in control neurons, from a mean phase of 28 degrees prior to the transition to 310 degrees immediately following (Watsons U test, p = 0.029; Fig. 5C and D). By comparison, neurons in MIA animals did not display a significant phase shift, with a mean phase of 261 degrees prior to the transition and 263 degrees afterwards (Watsons U test, p = 1; Fig. 5H and I). A comparison of control and MIA firing phase prior to the behavior transition revealed that they were significantly different (Watson U, p=0.028), however post-transition they were no longer so (Watson U, p=0.198).

**Figure 5.**
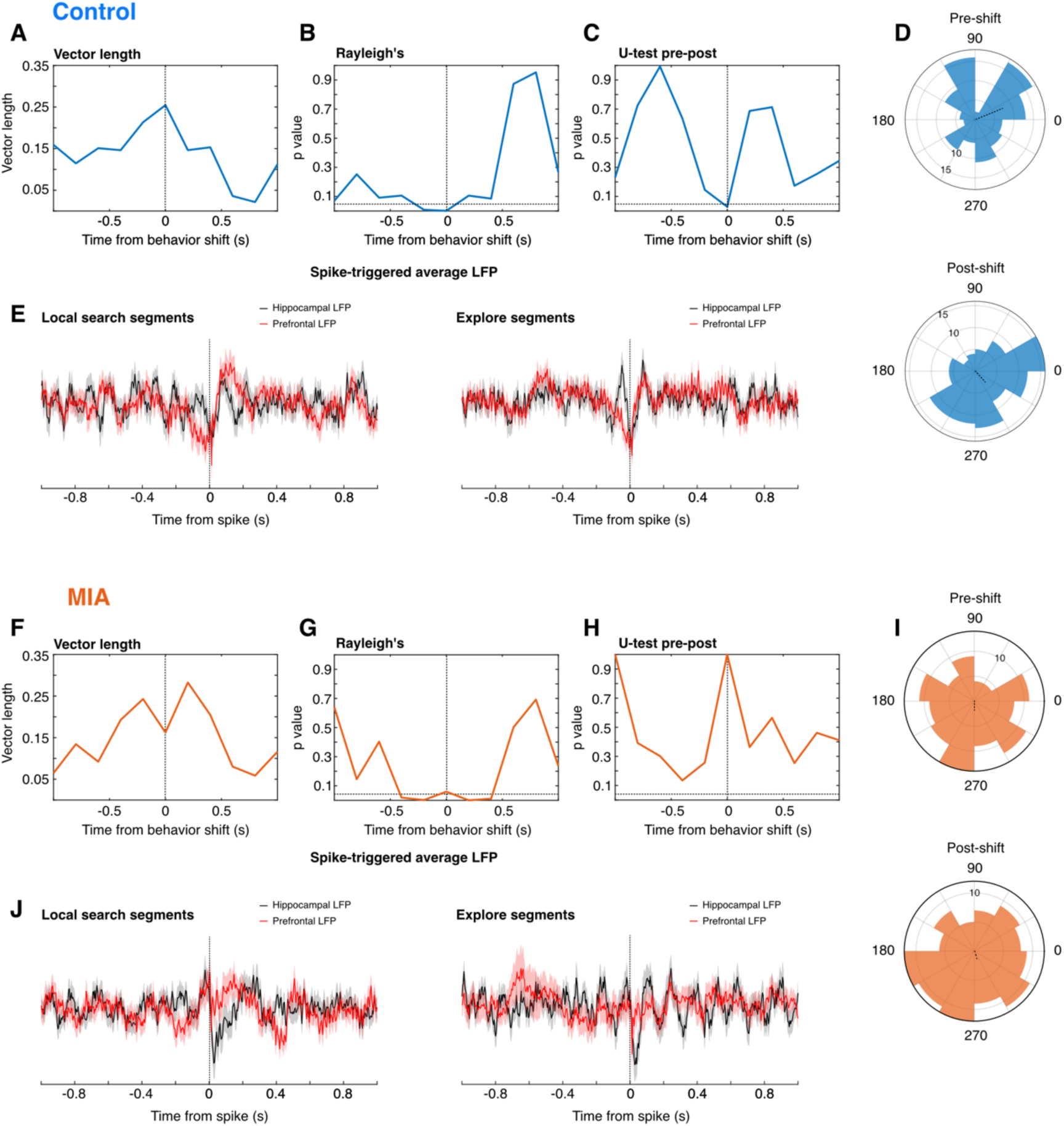
Single unit spike phases cluster during transitions. Firing phase of mPFC neurons, referenced to CA1 theta LFP, as control and MIA animals transition from exploration to local search. Firing becomes more entrained to theta following the behavioral transition, as indicted by an increase in vector length **(A,F)** and significant p values in the Rayleighs test **(B,G)**. A Watson’s U test indicates a significant shift in firing phase when 500ms of data pre and post transition is compared in control animals **(C)**, however, this change is not evident in MIA animals **(H)**. The change in firing phase pre-to-post behavior transition is also evident in the polar histogram of the preferred firing phase of neurons recorded in control animals **(D)**, but not in MIA animals **(I)**. **(E,J)** (E) shows 2 seconds of spike-triggered average LFP in hippocampus (black lines) and prefrontal cortex (red lines) ± SEM (shaded area). Left hand panels show spikes during the local search segments, while the right-hand panel shows explore segments. Vertical dotted lines show the time of the spike. (J) shows the same as (E), but for spikes and LFP from MIA animals.

### Partial Directed Coherence was greater during transition events

In control animals, across transition types, the primary direction of information flow was from hippocampus to prefrontal cortex (Figure 6A,B, Mean PDC ± SEM, Hippocampus to prefrontal = 0.52 ± 0.019, prefrontal to hippocampus = 0.33 ± 0.017, t(83) = 5.77, p < 0.0001). This was also true of the MIA animals (Figure 6F,G, Mean PDC ± SEM, Hippocampus to prefrontal = 0.45 ± 0.018, prefrontal to hippocampus = 0.32 ± 0.013, t(107) = 4.59, p < 0.0001). Compared to the control non-transition segments, there was significantly greater information flow from hippocampus to prefrontal and from prefrontal to hippocampus in both the search to explore, and explore to local search transitions (Figure 6D,C,E; Hippocampus to prefrontal, F(2,82) = 3.313, p = 0.0413; prefrontal to hippocampus, F(2,82) = 6.54, p = 0.0023). This was also true of the MIA animals from hippocampus to prefrontal, but not from prefrontal to hippocampus (Figure 6I,H,J; Hippocampus to prefrontal, F(2,106) = 8.16, p = 0.0005, prefrontal to hippocampus, F(2,106) = 2.59, p = 0.0794, n.s.). There was no difference in partial directed coherence between control and MIA animals in either transition condition (Figure 6K,L).

**Figure 6.**
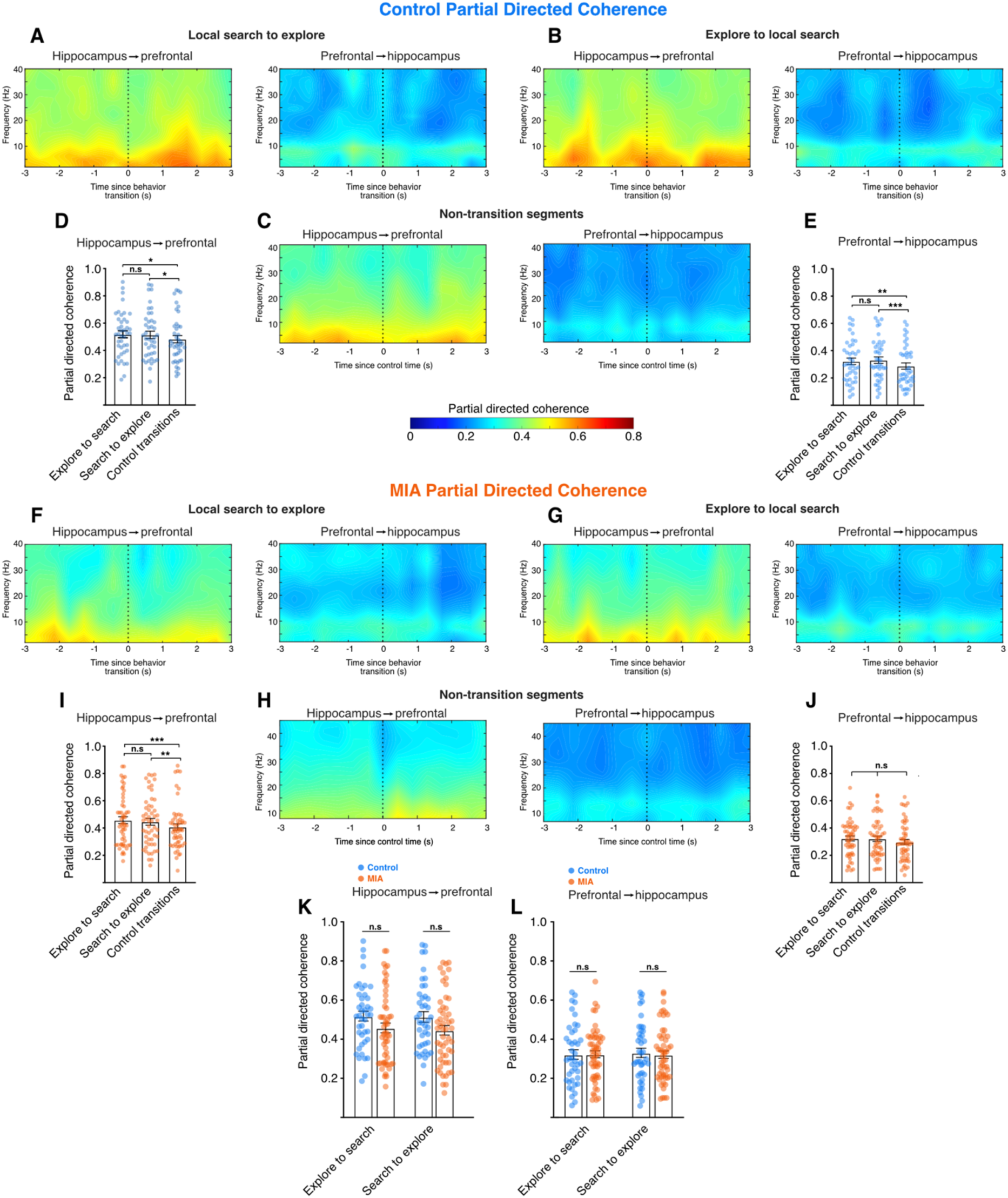
Information flows from hippocampus to prefrontal cortex during behavioral transitions. **(A,B,C)** Show the mean partial directed coherence from hippocampus to prefrontal cortex (left panels of each) and from prefrontal cortex to hippocampus (right panels of each). (A) shows local search to explore transitions (B) shows explore to search transitions and (C) shows control, non-transition segments. **(D,E)** The mean partial directed coherence (points) in each recording session for both transition types and control transitions, Solid bars show the mean, error bars illustrate the standard error. (D) shows data for hippocampus to prefrontal cortex, (E) shows prefrontal to hippocampus. **(F,G,H)** As for (A,B,C), but for sessions recorded from MIA animals. **(I,J)**, as for (D,E), but for sessions from MIA animals. **(K,L)** show the partial directed coherence in both transition types comparing control (blue) and MIA (orange) animals. (K) Shows partial directed coherence from hippocampus to prefrontal cortex, while (L) shows this data from prefrontal to hippocampus. (***p < 0.001, **p < 0.01, *p < 0.05).

## DISCUSSION

Our results show that the behavior of rats that are foraging freely in the open field is not uniform and can be algorithmically divided into at least two distinct modes of behavior, with regular transitions between them. The transitions we identify are between a high-speed, relatively linear behavior that moves the animal rapidly between locations, and a low-speed, highly tortuous and area-restricted behavior that is likely related to the animal making a decision to exploit a local region^40^ This finding is consistent with other studies that have investigated foraging strategies in rodents, showing that they switch between strategies, even in the absence of a specific task^41–43^. Furthermore, it indicates that studies using data obtained in the open field recording paradigm, as is commonly used in *in vivo* electrophysiological studies that reach as far back as the initial studies on hippocampal place cells^44–46^, should consider these more-nuanced behavior modes. Similarly, just as behavior in this task is not uniform, we describe clear peri-transition events in the local field potential of the hippocampus and prefrontal cortex that are not evident when animals are not at these behavioral transition points. In the hippocampus, there are clear differences in theta and beta power during behavioral transitions that have contrary depending on the direction of transition between the two types of behavior we identify. In the prefrontal cortex, these changes occur in the same way, but in the gamma band. Since hippocampal power in the theta band is associated with successful memory recall and encoding in rodents and humans^47,48^, it is possible that these decisions are related to spatial memory recall and encoding processes.

The hippocampus cooperates with prefrontal cortex in executive decision making^49–51^ and memory^52–54^, and we find that at the time of behavioral transition a burst of coherence occurs between prefrontal cortex and hippocampus in the theta band, as well as strong partial directed coherence data suggesting a strong flow of information from hippocampus to prefrontal cortex at these transition points that does not occur during non-transition times. Taken together, these findings suggest that as an animal makes a transition between behavioral modes, the hippocampus is involved in recalling information about where it is, or where it has been, and then communicating this to prefrontal cortex to guide the behavioral mode change.

The mechanisms that underlie the transition from local search to exploration and vice versa appear to be intact in the MIA animals, as the rhythmic transition from one strategy to the other is unchanged. More nuanced aspects of search patterns were, however, altered in MIA animals. MIA animals, tended to return to regions that had previously been the target of a local search, sooner than control animals. Furthermore, when search patterns were analyzed using a technique that treated local searches as if they were fixations across a visual image^35^, significant changes were also observed in the MIA animals. These animals made paths from one local search area to another that were more self-similar than those in controls, suggesting a tendency for stereotyped behavior. These differences were not due to changes in the quantity or duration of local search events or due to differences in thigmotaxis, as measures of these behaviors were similar in the two groups. It was also not due to variance in reward delivery, as this was randomized, and the same researcher ran both groups of animals. Rather, it supports the conclusion that the spatiotemporal structure of explore/exploit search patterns in MIA animals is different to that of controls, consistent with them having compromised memory for previously searched regions, possibly as a result of the changes in communication between hippocampus and prefrontal cortex observed in these animals.

When animals are foraging in areas where food is relatively evenly distributed across the environment, an adaptive behavior is to exploit the food locally (local search or area-restricted search) and then move to some more distal region (explore) before resuming exploitation. These exploration episodes have been characterized to occur as Lévy walks^17,55^, a type of random walk that creates clusters of local searching interspersed with relocations. While traditional levy walk models assume memoryless movement, incorporating a memory component can change patterns of activity. Here memory can either increase the probability that an animal will return to a productive patch (site fidelity) or alternatively, delay return until a site is replenished^56^. On this basis, one interpretation of our results is that in MIA animals there is a failure in the process by which memorial information about previous local search locations is communicated from the hippocampus to other brain regions, specifically, the prefrontal cortex. This is consistent with our findings that the directionality of this communication was primarily from hippocampus to prefrontal cortex during behavioural transitions, and that LFP coherence between the two structures is significantly reduced in MIA animals. Similarly, a shift in the phase of firing of prefrontal neurons relative to hippocampal LFP, that occurred during behavioural transitions in control animals, was absent in MIA animals. Previous evidence indicates that one role of prefrontal cortex in explore-exploit decisions is to encode subjective value and reward expectations^57^ and that the hippocampus has a role in learning to predict long-term reward^58^. Our results are, therefore, consistent with a model where memory for particular locations is linked to value via hippocampal input to prefrontal cortex, and that this communication is important for modulating exploit/explore decisions in the open field.

## REFERENCES

1. Stephens, D. W. & Krebs, J. R. Foraging Theory. vol. 6 (Princeton university press, 1986).

2. Hogeveen, J. et al. The neurocomputational bases of explore-exploit decision-making. Neuron 110, 1869–1879.e5 (2022).

3. Addicott, M. A., Pearson, J. M., Sweitzer, M. M., Barack, D. L. & Platt, M. L. A Primer on Foraging and the Explore/Exploit Trade-Off for Psychiatry Research. Neuropsychopharmacology 42, 1931–1939 (2017).

4. Cohen, J. D., McClure, S. M. & Yu, A. J. Should I stay or should I go? How the human brain manages the trade-off between exploitation and exploration. Philos. Trans. R. Soc. B Biol. Sci. 362, 933–942 (2007).

5. Costa, V. D., Mitz, A. R. & Averbeck, B. B. Subcortical Substrates of Explore-Exploit Decisions in Primates. Neuron 103, 533–545.e5 (2019).

6. Klein-Flügge, M. C., Bongioanni, A. & Rushworth, M. F. S. Medial and orbital frontal cortex in decision-making and flexible behavior. Neuron 110, 2743–2770 (2022).

7. Badre, D., Doll, B. B., Long, N. M. & Frank, M. J. Rostrolateral Prefrontal Cortex and Individual Differences in Uncertainty-Driven Exploration. Neuron 73, 595–607 (2012).

8. Daw, N. D., O’Doherty, J. P., Dayan, P., Seymour, B. & Dolan, R. J. Cortical substrates for exploratory decisions in humans. Nature 441, 876–879 (2006).

9. Eichenbaum, H. On the Integration of Space, Time, and Memory. Neuron 95, 1007– 1018 (2017).

10. O’Keefe, J. & Nadel, L. The Hippocampus as a Cognitive Map. (Clarendon Press; Oxford University Press, Oxford; New York, 1978).

11. Gauthier, J. L. & Tank, D. W. A Dedicated Population for Reward Coding in the Hippocampus. Neuron 99, 179–193.e7 (2018).

12. Sosa, M. & Giocomo, L. M. Navigating for reward. Nat. Rev. Neurosci. 22, 472–487 (2021).

13. Sosa, M., Plitt, M. H. & Giocomo, L. M. A flexible hippocampal population code for experience relative to reward. Nat. Neurosci. 1–13 (2025) doi:10.1038/s41593-025-01985-4.

14. Meck, W. H., Church, R. M. & Matell, M. S. Hippocampus, time, and memory—A retrospective analysis. Behav. Neurosci. 127, 642–654 (2013).

15. Dombrovski, A. Y., Luna, B. & Hallquist, M. N. Differential reinforcement encoding along the hippocampal long axis helps resolve the explore–exploit dilemma. Nat. Commun. 11, 5407 (2020).

16. Dorfman, A., Hills, T. T. & Scharf, I. A guide to area-restricted search: a foundational foraging behaviour. Biol. Rev. 97, 2076–2089 (2022).

17. Bartumeus, F., da Luz, M. G. E., Viswanathan, G. M. & Catalan, J. Animal Search Strategies: A Quantitative Random-Walk Analysis. Ecology 86, 3078–3087 (2005).

18. Estes, M. L. & McAllister, A. K. Maternal immune activation: Implications for neuropsychiatric disorders. Science 353, 772–777 (2016).

19. Sal-Sarria, S., Conejo, N. M. & González-Pardo, H. Maternal immune activation and its multifaceted effects on learning and memory in rodent offspring: A systematic review. Neurosci. Biobehav. Rev. 164, 105844 (2024).

20. Murray, B. G., Davies, D. A., Molder, J. J. & Howland, J. G. Maternal immune activation during pregnancy in rats impairs working memory capacity of the offspring. Neurobiol. Learn. Mem. 141, 150–156 (2017).

21. Baharnoori, M., Brake, W. G. & Srivastava, L. K. Prenatal immune challenge induces developmental changes in the morphology of pyramidal neurons of the prefrontal cortex and hippocampus in rats. Schizophr. Res. 107, 99–109 (2009).

22. Dickerson, D. D., Wolff, A. R. & Bilkey, D. K. Abnormal long-range neural synchrony in a maternal immune activation animal model of schizophrenia. J. Neurosci. Off. J. Soc. Neurosci. 30, 12424–31 (2010).

23. Wolff, A. R. & Bilkey, D. K. Prenatal immune activation alters hippocampal place cell firing characteristics in adult animals. Brain. Behav. Immun. 48, 232–243 (2015).

24. Munn, R. G. K., Wolff, A., Speers, L. J. & Bilkey, D. K. Disrupted hippocampal synchrony following maternal immune activation in a rat model. Hippocampus n/a, (2023).

25. Jing, Y., Zhang, H., Wolff, A. R., Bilkey, D. K. & Liu, P. Altered arginine metabolism in the hippocampus and prefrontal cortex of maternal immune activation rat offspring. Schizophr. Res. 148, 151–156 (2013).

26. Bilkey, D. K. & Muir, G. M. A low cost, high precision subminiature microdrive for extracellular unit recording in behaving animals. J. Neurosci. Methods 92, 87–90 (1999).

27. Zironi, I., Iacovelli, P., Aicardi, G., Liu, P. & Bilkey, D. K. Prefrontal Cortex Lesions Augment the Location-related Firing Properties of Area TE/Perirhinal Cortex Neurons in a Working Memory Task. Cereb. Cortex 11, 1093–1100 (2001).

28. Kyd, R. J. & Bilkey, D. K. Prefrontal cortex lesions modify the spatial properties of hippocampal place cells. Cereb. Cortex N. Y. N 1991 13, 444–51 (2003).

29. Kyd, R. J. & Bilkey, D. K. Hippocampal Place Cells Show Increased Sensitivity to Changes in the Local Environment Following Prefrontal Cortex Lesions. Cereb. Cortex 15, 720–731 (2005).

30. Muir, G. M. & Bilkey, D. K. Theta- and movement velocity-related firing of hippocampal neurons is disrupted by lesions centered on the perirhinal cortex. Hippocampus 13, 93–108 (2003).

31. Liu, P., Jarrard, L. E. & Bilkey, D. K. Excitotoxic lesions of the pre- and parasubiculum disrupt the place fields of hippocampal pyramidal cells. Hippocampus 14, 107–116 (2004).

32. Russell, N. A., Horii, A., Smith, P. F., Darlington, C. L. & Bilkey, D. K. Lesions of the Vestibular System Disrupt Hippocampal Theta Rhythm in the Rat. J. Neurophysiol. 96, 4–14 (2006).

33. Zar, J. H. Biostatistical Analysis: Pearson New International Edition. (2013).

34. Landler, L., Ruxton, G. D. & Malkemper, E. P. The multivariate analysis of variance as a powerful approach for circular data. Mov. Ecol. 10, 21 (2022).

35. Cristino, F., Mathôt, S., Theeuwes, J. & Gilchrist, I. D. ScanMatch: A novel method for comparing fixation sequences. Behav. Res. Methods 42, 692–700 (2010).

36. Anderson, N. C., Anderson, F., Kingstone, A. & Bischof, W. F. A comparison of scanpath comparison methods. Behav. Res. Methods 47, 1377–1392 (2015).

37. Bella-Fernández, M., Suero Suñé, M. & Gil-Gómez de Liaño, B. Foraging behavior in visual search: A review of theoretical and mathematical models in humans and animals. Psychol. Res. 86, 331–349 (2022).

38. Omidvarnia, A., Mesbah, M., O’Toole, J. M., Colditz, P. & Boashash, B. Analysis of the time-varying cortical neural connectivity in the newborn EEG: A time-frequency approach. in International Workshop on Systems, Signal Processing and their Applications, WOSSPA 179–182 (2011). doi:10.1109/WOSSPA.2011.5931445.

39. Berens, P. CircStat: a MATLAB toolbox for circular statistics. (2009) 10.18637/jss.v031.i10.

40. Jackson, B. J., Fatima, G. L., Oh, S. & Gire, D. H. Many Paths to the Same Goal: Balancing Exploration and Exploitation during Probabilistic Route Planning. eNeuro 7, (2020).

41. Gire, D. H., Kapoor, V., Arrighi-Allisan, A., Seminara, A. & Murthy, V. N. Mice develop efficient strategies for foraging and navigation using complex natural stimuli. Curr. Biol. CB 26, 1261–1273 (2016).

42. Neville, V. et al. You are How You Eat: Foraging Behavior as a Potential Novel Marker of Rat Affective State. Affect. Sci. 5, 232–245 (2024).

43. Inglis, I. R. et al. Foraging behaviour of wild rats (*Rattus norvegicus*) towards new foods and bait containers. Appl. Anim. Behav. Sci. 47, 175–190 (1996).

44. O’Keefe, J. & Dostrovsky, J. The hippocampus as a spatial map. Preliminary evidence from unit activity in the freely-moving rat. Brain Res. 34, 171–175 (1971).

45. Muller, R. & Kubie, J. The firing of hippocampal place cells predicts the future position of freely moving rats. J. Neurosci. Off. J. Soc. Neurosci. 9, 4101–10 (1989).

46. Thompson, L. & Best, P. Place cells and silent cells in the hippocampus of freely-behaving rats. J. Neurosci. Off. J. Soc. Neurosci. 9, 2382–90 (1989).

47. Kota, S., Rugg, M. D. & Lega, B. C. Hippocampal Theta Oscillations Support Successful Associative Memory Formation. J. Neurosci. 40, 9507–9518 (2020).

48. Joensen, B. H. et al. Hippocampal theta activity during encoding promotes subsequent associative memory in humans. Cereb. Cortex N. Y. N 1991 33, 8792– 8802 (2023).

49. Bai, W. et al. Hippocampal-prefrontal high-gamma flow during performance of a spatial working memory. Brain Res. Bull. 207, 110887 (2024).

50. Yu, J. Y. & Frank, L. M. Hippocampal–cortical interaction in decision making. Neurobiol. Learn. Mem. 117, 34–41 (2015).

51. Gluth, S., Sommer, T., Rieskamp, J. & Büchel, C. Effective Connectivity between Hippocampus and Ventromedial Prefrontal Cortex Controls Preferential Choices from Memory. Neuron 86, 1078–1090 (2015).

52. Cohen, M. X. Hippocampal-Prefrontal Connectivity Predicts Midfrontal Oscillations and Long-Term Memory Performance. Curr. Biol. 21, 1900–1905 (2011).

53. Backus, A. R., Schoffelen, J.-M., Szebényi, S., Hanslmayr, S. & Doeller, C. F. Hippocampal-Prefrontal Theta Oscillations Support Memory Integration. Curr. Biol. 26, 450–457 (2016).

54. Guise, K. G. & Shapiro, M. L. Medial Prefrontal Cortex Reduces Memory Interference by Modifying Hippocampal Encoding. Neuron 94, 183–192.e8 (2017).

55. de Jager, M., Weissing, F. J., Herman, P. M. J., Nolet, B. A. & van de Koppel, J. Lévy Walks Evolve Through Interaction Between Movement and Environmental Complexity. Science 332, 1551–1553 (2011).

56. Kazimierski, L. D., Abramson, G. & Kuperman, M. N. Random-walk model to study cycles emerging from the exploration-exploitation trade-off. *Phys*. Rev. E 91, 012124 (2015).

57. Padoa-Schioppa, C. & Assad, J. A. Neurons in the orbitofrontal cortex encode economic value. Nature 441, 223–226 (2006).

58. Stachenfeld, K. L., Botvinick, M. M. & Gershman, S. J. The hippocampus as a predictive map. Nat. Neurosci. 20, 1643–1653 (2017).

